# Disease modeling by efficient genome editing using a near PAM-less base editor *in vivo*

**DOI:** 10.1101/2021.06.28.450169

**Authors:** Marion Rosello, Malo Serafini, Marina C Mione, Jean-Paul Concordet, Filippo Del Bene

**Affiliations:** Sorbonne Université, INSERM U968, CNRS UMR 7210, Institut de la Vision, Paris, France; Department of Cellular, Computational and Integrative Biology – CIBIO, University of Trento, Trento, Italy; Muséum National d’Histoire Naturelle, INSERM U1154, CNRS UMR 7196, Paris, France

## Abstract

Base Editors are emerging as an innovative technology to introduce point mutations in complex genomes. So far, the requirement of an NGG Protospacer Adjacent Motif (PAM) at a suitable position often limits the editing possibility to model human pathological mutations in animals. Here we show that, using the CBE4max-SpRY variant recognizing the NRN PAM sequence, we could introduce point mutations for the first time in an animal model and achieved up to 100% efficiency, thus drastically increasing the base editing possibilities. With this near PAM-less base editor we could simultaneously mutate several genes and developed a co-selection method to identify the most edited embryos based on a simple visual screening. Finally, we applied our method to create a new zebrafish model for melanoma predisposition based on the simultaneous editing of multiple genes. Altogether, our results considerably expand the Base Editor application to introduce human disease-causing mutations in zebrafish.

## Introduction

The CRISPR (Clustered Regularly Interspaced Short Palindromic Repeats)/Cas9 system is a very powerful gene editing tool to perform mutagenesis in zebrafish^1^. The sgRNA-Cas9 complex first binds to target DNA through a Protospacer Adjacent Motif (PAM) and Cas9 next cleaves DNA upon stable annealing of the sgRNA spacer sequence to the sequence immediately upstream of the PAM. The NGG PAM sequence is thus required for the *Sp*Cas9 protein to introduce a DNA double-strand break (DSB). This technique is now extensively used in zebrafish to produce knock-out alleles^2^. Additionally, recent studies showed that exogenous DNA and Single Nucleotide Polymorphisms (SNPs) can be introduced in the genome using CRISPR/Cas9 mediated homology-directed (HDR) repair with variable efficiency^3, 4, 5^. In order to complement these HDR-based strategies to introduce specific point mutation, second-generation gene editing tools called base editors (BEs) have recently been developed. The Cytidine Base Editor (CBE) is composed of a Cas9-D10A nickase fused to a cytidine deaminase and converts C-to-T bases in a restricted window of 13 to 19 nucleotides (nt) upstream of the PAM sequence without generating DSBs^6, 7, 8^. In zebrafish, several CBE variants have been shown to work with different gene editing efficiencies^9, 10, 11, 12^. In a previous study we tested several CBE variants and we were able to reach a C-to-T conversion efficacy of 91% in zebrafish, showing many applications from signaling pathway activation to human disease modeling^12^. It has been shown that potentially all C bases within the PAM [-19, -13] bp window can be edited with these tools, although a higher efficiency was generally achieved for the Cs located in the middle of this editing window, highlighting the importance of the C distance to the PAM for efficient base editing^13^. Therefore, the restriction of the base modification to the PAM-dependent window is still a critical intrinsic limitation to the base editor techniques. Due to this restriction, these tools cannot be applied to any gene and any mutation of interest. To overcome this limitation, extensive works have been done in cultured cells to engineer CBEs recognizing other PAM sequences than the classical NGG and thereby significantly expand the range of C bases that can be converted. Based on the technological advances made in cultured cells, we tested several novel CBEs in zebrafish to overcome the PAM limitation. Among them, the most recent and flexible variant is the CBE4max-SpRY, reported as a near PAM-less base editor recognizing almost all PAM sequences in cultured cells^14^. Here we established the CBE4max-SpRY CBE variant for the first time in an animal model. Using this variant, we were able to perform in zebrafish C-to-T conversions at an unprecedented high efficiency, reaching up to 100%. We show that the CBE4max-SpRY converts C bases efficiently using NG or NA PAM in zebrafish and that it is possible to mutate several genes using NG and NA PAMs at the same time, increasing drastically the base editing possibilities. Based on these results, we developed a co-selection method to phenotypically identify the most edited embryos following the CBE4max-SpRY, by co-targeting for the *tyrosinase* gene and selecting embryos for lack of pigmentation. Finally, using this approach, we could for the first time in zebrafish simultaneously introduce a loss-of-function mutation in a tumor suppressor gene together with a gain-of-function mutation in an endogenous oncogene. We targeted *tp53* tumor suppressor and *nras* oncogene and generated a new zebrafish model with an abnormal melanocyte growth which is an hallmark of melanoma formation susceptibility^15^, without over-expressing mutated oncogenes which has been the main strategy used in zebrafish to model cancer so far^16, 17, 18, 19^.

## Results

### Evaluation of several CBE variants recognizing non-NGG PAM sequences in zebrafish

Base editing requires the presence of a PAM at 13 to 19 bp downstream of the targeted C base. This limitation is critical and often makes CBEs unable to introduce the desired point mutation in the genome. To overcome this limitation particularly important when trying to model disease causing mutations in animal models, we first tested three different CBE variants recognizing the NG PAM: xCas9-BE4^20^, CBE4max-SpG^14^ and CBE4max-SpRY^14^. In order to analyze their efficacy in zebrafish, we started by injecting into one-cell stage embryos sgRNAs acting upstream of an NGG PAM that we previously found to be very efficient with the original CBEs^12^. First we tested *tp53 Q170\** sgRNA, with which we got up to 86% of efficiency using BE4-gam^12^. After injection with the *xCas9-BE4* mRNA, we were unable to detect any base conversion by Sanger sequencing of PCR products of the target region. We then tested CBE4max-SpG and CBE4max-SpRY with *rb1W63*NGG* sgRNA targeting the *rb1* tumor suppressor gene upstream of an NGG PAM (Fig. 1a) to introduce the W63* mutation with which we previously got up to 91% of base conversion using the ancBE4max variant^12^. We sequenced the target region from 14 single embryos and we did not detect any C-to-T conversion using CBE4max-SpG whereas we got C-to-T conversions for 4 out of 11 single embryos analyzed with up to 33% of efficiency using CBE4max-SpRY (Fig. 1b). The CBE4max-SpRY has been reported as a near PAM-less base editor in cultured cells^14^ and we next chose to analyze its flexibility. We designed two sgRNAs, one upstream of an NG PAM (*rb1W63*NG* sgRNA) and the second upstream of an NA PAM (*rb1W63*NA* sgRNA), in order to introduce the same mutation in the *retinoblastoma1* (*rb1*) gene (Fig. 1a). After injection into one-cell stage of each sgRNAs with the *CBE4max-SpRY* mRNA, we sequenced two pools of 9 injected embryos. With both sgRNAs we obtained high base editing efficiency rates, up to 58% of efficiency with the NG PAM and up to 48% of efficiency with the NA PAM (Fig. 1c). Additionally, we noticed that another C was also edited, leading to the R64K mutation but it was 3’ to the premature STOP codon introduced (C16 for the NG sgRNA and C17 for the NA sgRNA). In our previous study using the ancBE4max CBE, this C was at 19 bp from the NGG PAM and we obtained less than 22% of base editing^12^ whereas here we show that this C can be converted up to 55% of efficiency (Fig. 1c). This last result shows that for some C base targets, thanks to the PAM flexibility of the CBE4-SpRY, we can now increase the C-to-T conversion efficacy compared to the use of the classical ancBE4max. We demonstrated here for the first time in animals that the near PAM-less CBE4max-SpRY converts a C base into a T base efficiently using NA or NG PAMs.

**Figure 1.**
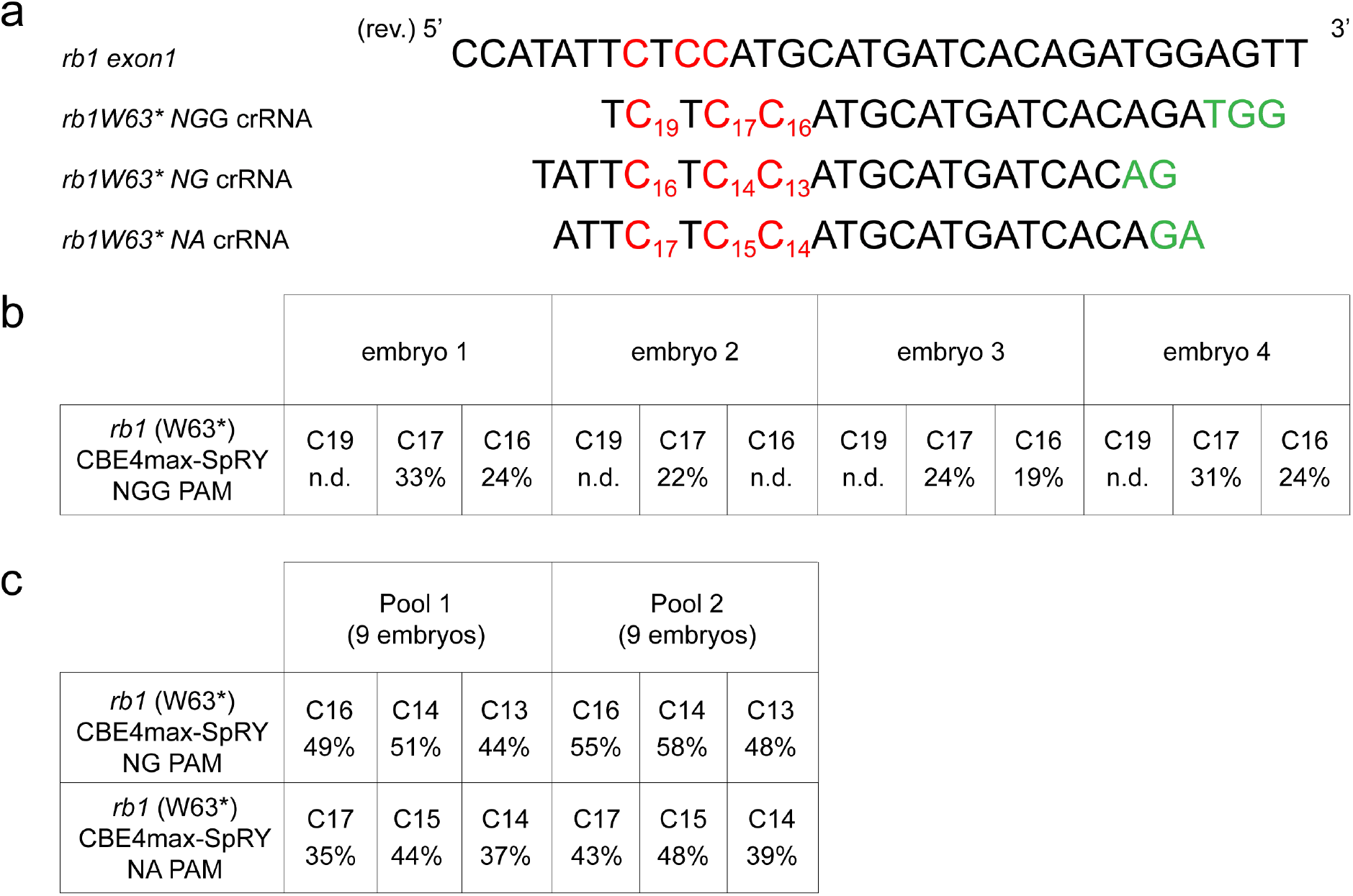
Efficient C-to-T conversion using the CBE4max-SpRY variant with NA and NG PAMs. **a**. Targeted genomic sequence of the exon 1 of the *rb1* tumor suppressor gene and the 3 different sgRNAs used to introduce the W63* mutation. For each sgRNA, the targeted Cs are in red and the PAM sequence is in green. **b**. C-to-T conversion efficiency for each targeted Cs in the 4 single edited embryos out of 11 analyzed embryos injected with *CBE4max-SpRY* mRNA and *rb1W63* NGG* sgRNA. **c**. C-to-T conversion efficiency for each targeted Cs in 2 pools of 9 embryos injected with *CBE4max-SpRY* mRNA and *rb1W63* NGG* sgRNA.

### Total base conversion in F0 embryos using an NA PAM

In order to explore the efficiency of the near PAM-less CBE variant in zebrafish, we decided to target another locus, the *tyrosinase* gene. We designed 3 different guides upstream, respectively, of an NGG, NA and NG PAM in order to introduce the W273* mutation in Tyrosinase, an enzyme necessary for the production of melanin pigments (Fig. 2a). Upon independent injections of these sgRNAs with the *ancBE4max* mRNA for the NGG sgRNA and the C*BE4max-SpRY* mRNA for the NGG, NA and NG sgRNAs, the embryos showed a range of pigmentation defects. We divided these phenotypes in four groups depending on the severity of the pigmentation defects: wild-type like, mildly depigmented, severely depigmented and albino. Representative pictures of 2 days post-fertilization (2 dpf) embryos for each group are illustrated in Fig. 2c. Upon the injection of the *tyrW273* NGG* sgRNA and the *ancBE4max* mRNA, we obtained 50% of mildly depigmented embryos with a small proportion of severely affected embryos whereas surprisingly almost 100% of the injected fish with the *CBE4max-SpRY* mRNA were depigmented (Fig. 2b, columns 2 and 3). Next, using the *tyrW273* NA* sgRNA and the CBE4max-SpRY, we obtained 100% of injected embryos showing pigmentation defects with almost 50% exhibiting a total lack of pigmentation (Fig. 2b, column 4). Finally, with the NG PAM sgRNA, 50% of the injected embryos were poorly affected, the base editing efficiency reached only 20% (Fig. 2b columns 5, d). Remarkably in the pool of 9 albino embryos from the injection of the *tyrW273* NA* sgRNA and *CBE4max-SpRY* mRNA, we obtained 100% of C-to-T conversion for the C16 and 95% for the C15 (Fig. 2d, e), an efficiency rate never reached so far even with the use of the classical CBEs recognizing the NGG PAM. We did not obtain this C-to-T efficiency neither using the *tyrW273* NG* sgRNA nor the ancBE4max with the *tyrW273* NGG* sgRNA (Fig. 2b, d). We could speculate that the difference of base conversion efficiency is due to the distance of the C from the PAM or to the sgRNA sequence for which the shift of one base would drastically impact the gene editing efficiency (Fig. 2a). Moreover, we found that the mutation was transmitted to 100% of the offspring by screening only 4 F0 adult fish presenting pigmentation defect at the adult stage (Supplementary Fig. 1, Founder 1). Other 3 F0 fish also transmitted the mutation (Supplementary Fig. 1). The screening was performed by Sanger sequencing of PCR products of the *tyrosinase* region in random single F1 embryos from an outcross of each founder. Moreover, by crossing 2 founders, Founder 1 and 2 in supplementary Fig. 1, we were able to obtain 51,9% of albino embryos (n=28/54). Together these results highlight the CBE4max-SpRY as a very powerful CBE variant, showing that for many targets this CBE could be a better choice than the classical CBEs and could edit 100% of the alleles of the injected embryo using NA or NG PAMs.

**Figure 2.**
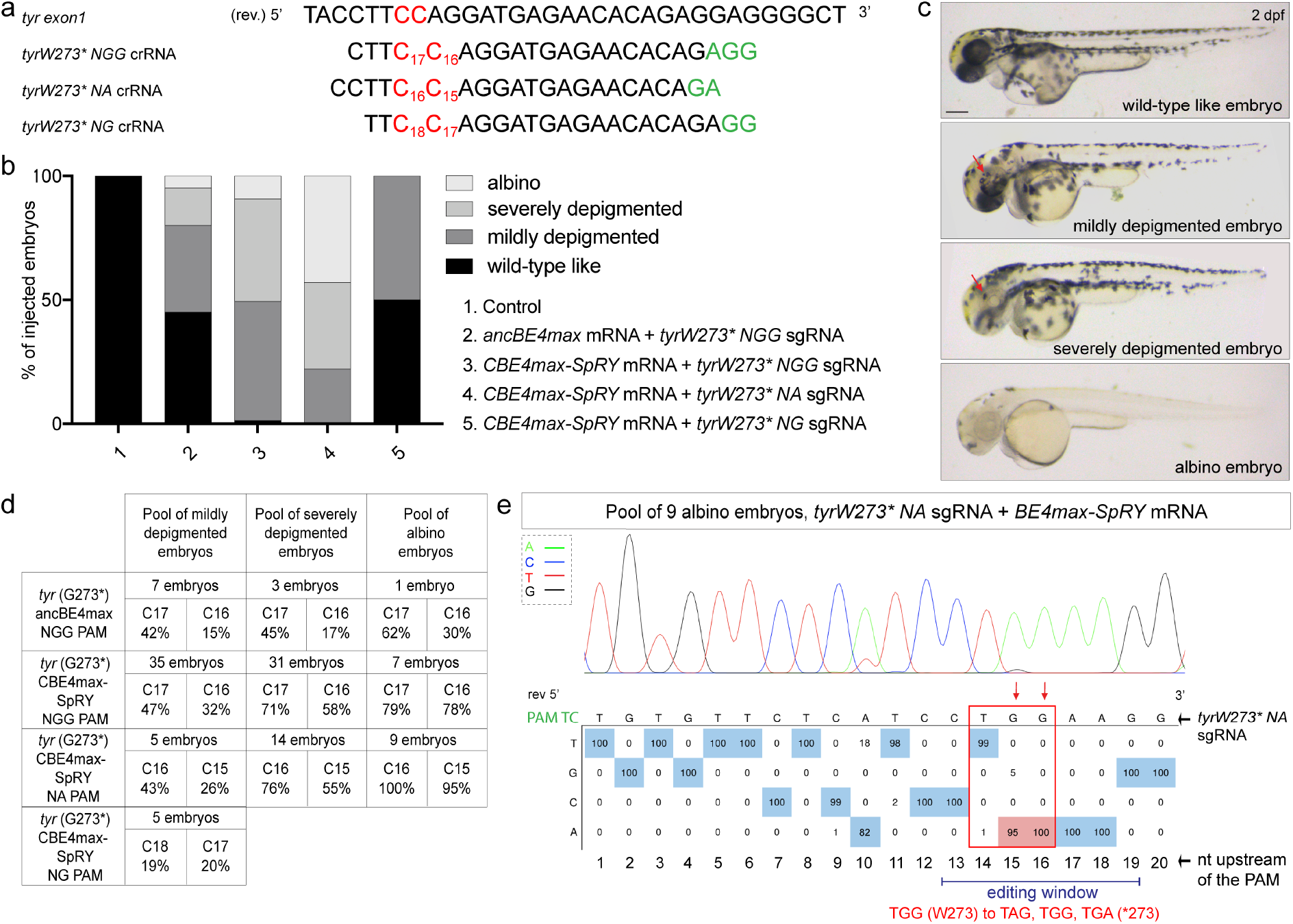
Total base conversion in the F0 embryos with CBE4max-SpRY and a NA PAM. **a**. Targeted genomic sequence of the exon 1 of the *tyrosinase* gene and the 3 different sgRNAs used to introduce the W273* mutation. For each sgRNA, the targeted Cs are in red and the PAM sequence is in green. **b**. Proportion of the 4 groups based on the pigmentation defects described in Fig. 2c for each injection: the *ancBE4max* mRNA and the *tyrW273*NGG* sgRNA (column 2, 19 embryos in total), the *CBE4max-SpRY* mRNA and the *tyrW273*NGG* sgRNA (column 3, 74 embryos in total), the *CBE4max-SpRY* mRNA and the *tyrW273*NA* sgRNA (column 4, 28 embryos in total), the *CBE4max-SpRY* mRNA and the *tyrW273*NG* sgRNA (column 5, 10 embryos in total). **c**. Lateral view of representative 2 days post-fertilization (dpf) embryos showing different severity of pigmentation defects (wild-type like, mildly depigmented, severely depigmented and albino embryos). Scale bar = 100 µm. **d**. C-to-T conversion efficiency for each targeted Cs and each pool of embryos showing pigmentation defects presented in Fig. 2b. **e**. DNA sequencing chromatogram of the targeted *tyr* gene from a pool of 9 albino embryos obtained after the injection of the *CBE4max-SpRY* mRNA with the *tyrW273*NA* sgRNA. W273* mutation in Tyr upon C-to-T conversion in *tyr* reached 100% for the C16 base and 95% for the C15 base of gene editing efficiency. Numbers in the boxes represent the percentage of each base at that sequence position. In red are highlighted the base substitutions introduced by base editing while the original bases are in blue. The color code of the chromatogram is indicated in the upper left corner (Adenine green, Cytosine blue, Thymine red, Guanine black). The distance from the PAM sequence of the targeted C base is indicated below the chromatogram^31^.

### Simultaneous editing of two different bases

To test if we can take advantage to the high flexibility and efficacy of the CBE4max-SpRY variant in zebrafish for multiplex editing, we targeted two loci at the same time using a sgRNA upstream of an NG PAM and one upstream of an NA PAM. We injected the *CBE4max-SpRY* mRNA and the two synthetic *tyrW273* NA* and *rb1W63* NG* sgRNAs into the cell at one-cell stage. Among the injected embryos, 100% showed pigmentation defects and at least 50% exhibited total absence of pigmentation (Fig. 3a). It can be noted that this proportion is almost the same as the one after the use of the *tyrW273* NA* sgRNA only and the *CBE4max-SpRY* mRNA (Fig. 2b, column 4), meaning that the addition of a second guide did not affect the base conversion efficacy at the *tyrosinase* locus. Sequencing of the two loci was performed on three different pools of embryos separated according to the severity of their pigmentation defects. As expected, the severity of the pigmentation phenotype follows the base conversion efficiency of the *tyrosinase* gene (Fig. 3a, b). In particular, we found that in the pool of 35 albino embryos, we could no longer detect the wild type C by Sanger sequencing, suggesting almost 100% of C-to-T conversion in the *tyrosinase* gene. Remarkably, in the albino pool, up to 44% of C-to-T conversion of the *rb1* gene was detected, revealing that double *tyrosinase* and *rb1* mutations were present in a large proportion of cells (Fig. 3a, b), while in mildly depigmented zebrafish, editing of *rb1* was up to 16%. In addition, we found that both mutations were transmitted to the offspring with high transmission rates, as shown by screening the progeny of only one F0 adult fish (Fig. 3c, d). We obtained different combinations of mutations, and we showed that some F1 embryos were mutated for the *tyrosinase* gene alone. At the same time, we noticed that some embryos were mutated for different Cs of the editing windows for each gene (Fig. 3d). With these results we could demonstrate that, using this approach, it is now possible to mutate simultaneously two different genes with high efficiency by combining two different PAM sequences in zebrafish, NA and NG PAMs. Additionally, we can observe that the highest efficiency for *rb1* mutation was found in the albino embryo pool, embryos for which the *tyr* mutation is generated in 100% of the alleles as measured by Sanger sequencing.

**Figure 3.**
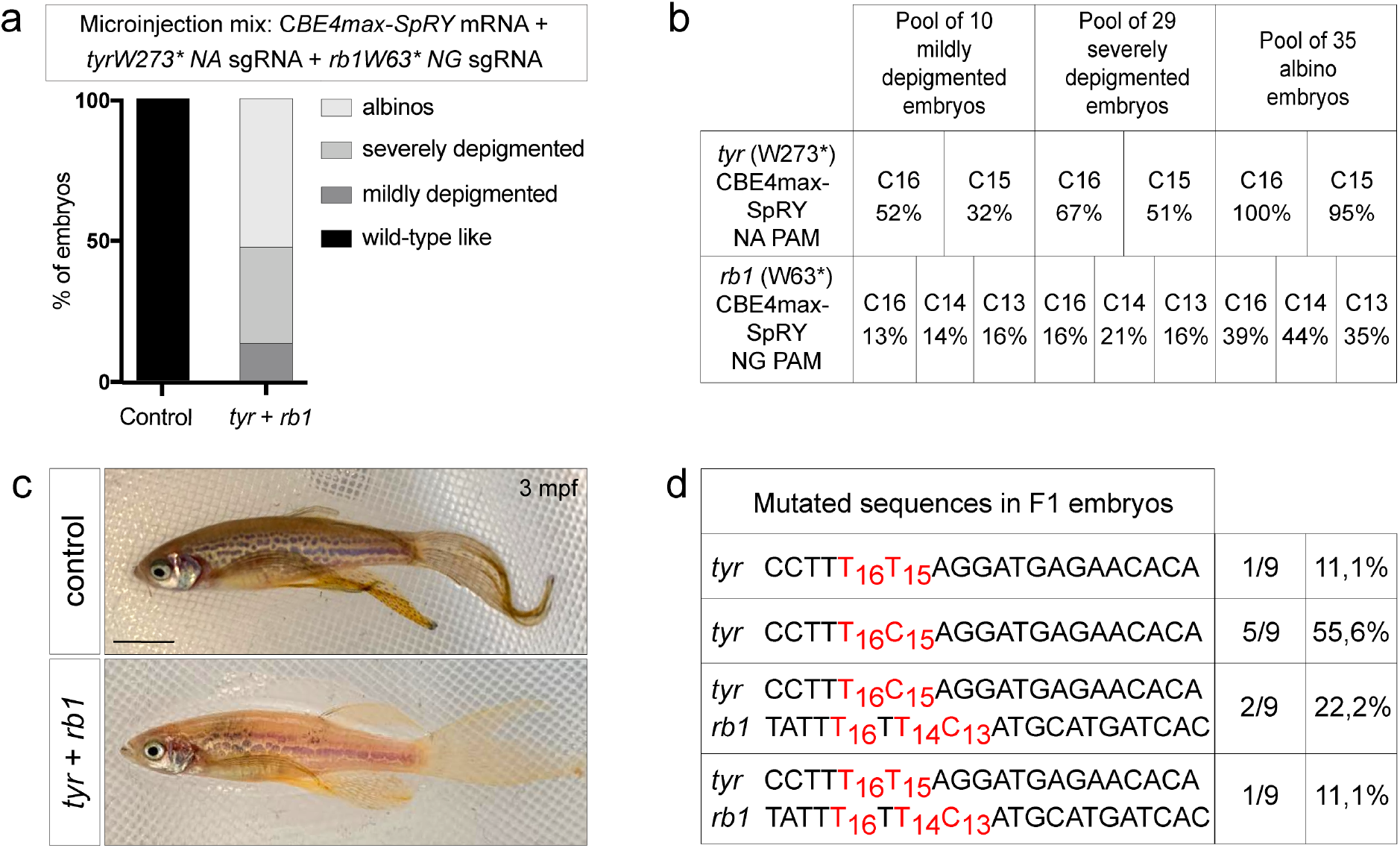
Germline transmission of two mutations generated simultaneously using NA and NG PAMs. **a**. Proportion of the 4 groups based on the pigmentation lack defects described in Fig. 2c for embryos injected with the *CBE4max-SpRY* mRNA, the *tyrW273*NA* and the *rb1W63*NG* sgRNA (column 2, 74 embryos in total). **b**. C-to-T conversion efficiency for each targeted Cs and each pool of embryos showing pigmentation defects presented in Fig. 3a for the *tyrosinase* and *rb1* genes. **c**. Lateral view of a 3 months F0 fish injected at one-cell stage embryo with the *CBE4max-SpRY* mRNA, the *tyrW273*NA* and the *rb1W63*NG* sgRNAs and showing pigmentation defects. Scale bar = 5 mm. **d**. Sequenced *tyr* and *rb1* loci of 9 F1 single embryos from the founder fish in Fig. 3c. 6 embryos out of 9 were edited for *tyrosinase* and 3 embryos out of 9 were double edited for *tyrosinase* and *rb1*.

### Co-selection strategy to prescreen phenotypically the most edited embryos

The efficiency of CBE4max-SpRY achieved here above opens the possibility to perform multiplex mutagenesis in zebrafish. Such experiments, however, usually involve time consuming screening to obtain a founder carrying all the desired mutations or long crossing protocols to genetically combine multiple mutations identified in different animals. For this reason, we have decided to take advantage of the high base editing rate of the *tyrW273*NA* sgRNA and developed a method for co-selection of base editing to rapidly identify the most edited F0 embryos following *CBE4max-SpRY* mRNA injections. We first injected the *rb1W63*NG* and *nras NA* sgRNAs with the near PAM-less *CBE4max-SpRY* mRNA and found C-to-T conversion rates up to 3% at *rb1* and 50% at *nras* targets in a pool of 50 injected embryos (Fig. 4a). Moreover, we can observe that the C12 has been edited at 23% of efficiency and the C18 has not been edited on the contrary of what we usually observe using the other CBE4 variants (Fig. 4a). From these observations, we can speculate that the editing windows of the CBE4max-SpRY is slightly different than the usual [-19, -13 bp] PAM window and that with the use of this CBE it is possible to target the C12 bases. We then added to the same micro-injection mix the *tyrW273*NA* sgRNA and obtained embryos showing pigmentation defects that we split in four groups as above (Fig. 2c and Fig. 4b). By sequencing the 3 loci in each pool of embryos, we could show that the editing efficiency is higher or lower to the same extent in the 3 targeted loci (Fig. 4c). The mildly depigmented embryos were edited up to 41% for *tyr*, 61% for *nras* and no detectable mutation for *rb1* whereas the albino embryos were edited up to 95% for *tyr*, 100% for *nras* and 22% for *rb1* (Fig. 4c). Base editing co-selection strategies were recently demonstrated in cultured cells^21^ and have not been reported in animals so far. Using the *tyrW273* NA* sgRNA, we show here a powerful strategy to readily obtain the most efficiently C base edited embryos at targeted loci of interest by simply selecting for albino embryos resulting from co-targeting the *tyrosinase* gene.

**Figure 4.**
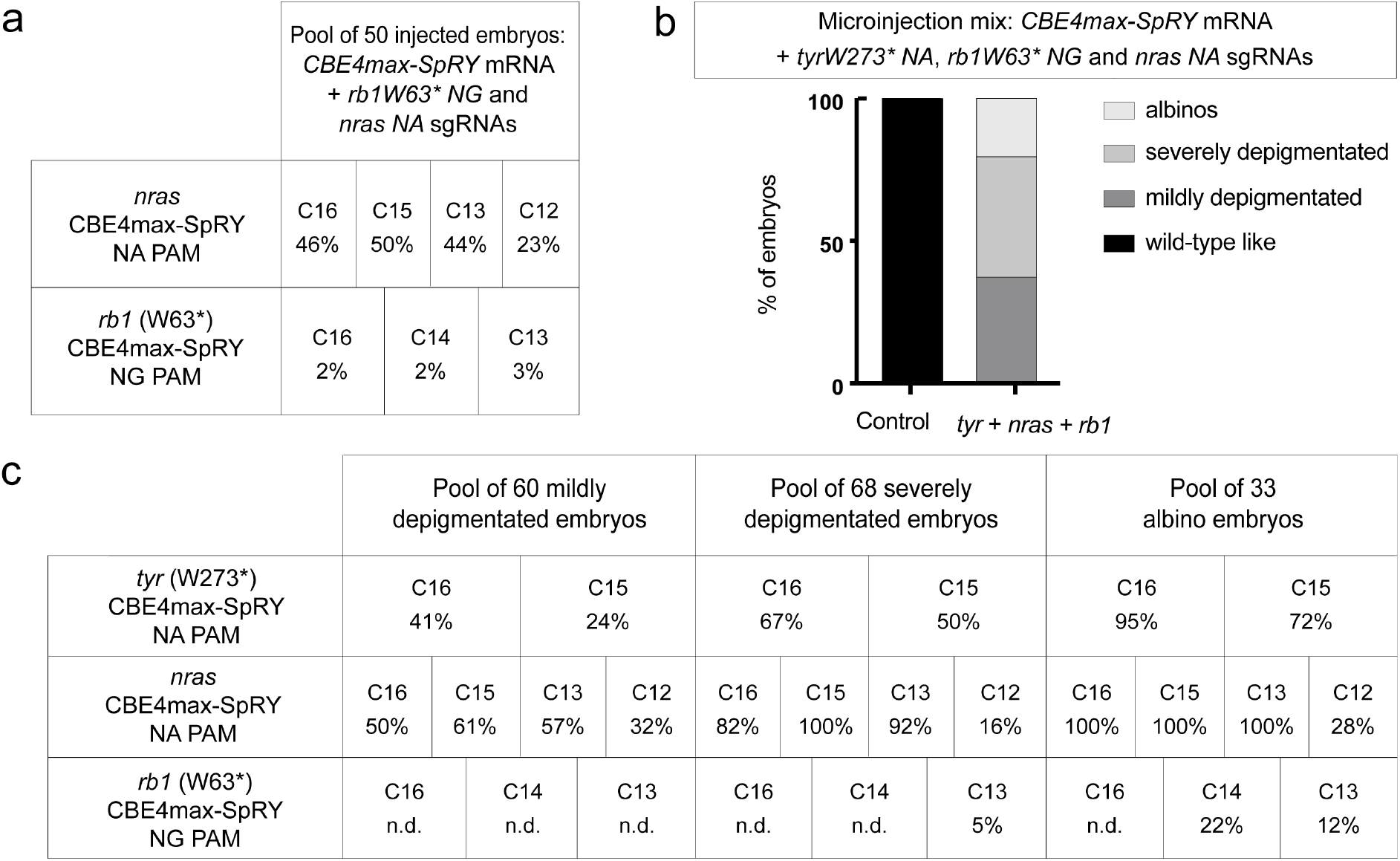
Base editing co-selection method. **a**. C-to-T conversion efficiency for each targeted Cs in *nras* and *rb1* genes in a pool of 50 embryos injected with *CBE4max-SpRY* mRNA and *nras NA* and *rb1 W63\** sgRNAs. **b**. Proportion of the 4 groups based on the pigmentation defects described in Fig. 2c for the embryos injected with the *CBE4max-SpRY* mRNA, the *tyrW273*NA, nras NAA* and *rb1W63*NG* sgRNAs (column 2, 161 embryos in total). **c**. C-to-T conversion efficiency for each targeted Cs and each pool of embryos showing pigmentation defects presented in Fig. 4b for the *tyrosinase, nras* and *rb1* genes. The albino embryos are the most edited for the 3 different loci.

### Genetic disease modeling by generating combination of mutations

We next investigated the ability to use CBE4max-SpRY to introduce in zebrafish mutations found in human cancers. The zebrafish has become a powerful *in vivo* model to study a high variety of human cancer types^16, 17, 18, 19^. However, studies to assess the activity of mutations in oncogenes in this animal model have relied so far on transgenic approaches that express human oncogenes using tissue specific or constitutive promoters. In order to more accurately mimic the impact of cancer mutations as observed in somatic alterations in humans, we decided to use the CBE4max-SpRY to directly generate combination of mutations in endogenous zebrafish genes preserving their normal transcriptional regulation. To test this important innovative approach, we aimed at generating cancer models with Nras oncogene activation combined with loss-of-function mutations in the *tp53* tumor suppressor gene. We thus injected into one-cell stage embryos the *CBE4max-SpRY* mRNA, *nras NAA* and *tp53 Q170\** sgRNAs. Upon this injection we found 50% of the injected fish with an over-all increase of pigmentation at 3 dpf (n=36/70), a phenotype never seen upon separate injection of each sgRNA alone (Fig. 5a). To verify that the absence of phenotype when injecting only *nras NAA* sgRNA is not due to a low efficiency of gene editing, we sequenced a pool of 80 injected embryos and we detected high efficiency of base conversion (Fig. b). We next randomly selected 4 embryos presenting a pigmentation similar to control embryos and 8 embryos showing an increase of pigmentation after the injection of the sgRNAs targeting *nras* and *tp53* genes. After sequencing both loci, we found indeed a correlation between the pigment phenotype and the multi-mutagenesis efficiency, embryos with an increase of pigmentation were more edited for *nras* and *tp53* genes (Fig. 5c, d). We thus have shown for the first time that endogenous activation of *nras* oncogene and knock-out of *tp53* tumor suppressor gene leads to an increase of melanocyte numbers in zebrafish, an early evidence of abnormal melanocyte growth which could led to melanoma formation (Fig. 5). Indeed, it has been reported that fish over-expressing human mutated HRAS oncogene in melanocytes were hyperpigmented at 3 dpf and developed melanoma at the adult stage^15^. Moreover, other reports using zebrafish transgenic lines have suggested a role of p53 and Ras oncogenes in melanoma formation^22, 23^. We have developed here for the first time a hyper-pigmentation zebrafish model by generating endogenous activating mutation in *nras* oncogene and loss-of-function mutation in *tp53* tumor suppressor gene.

**Figure 5.**
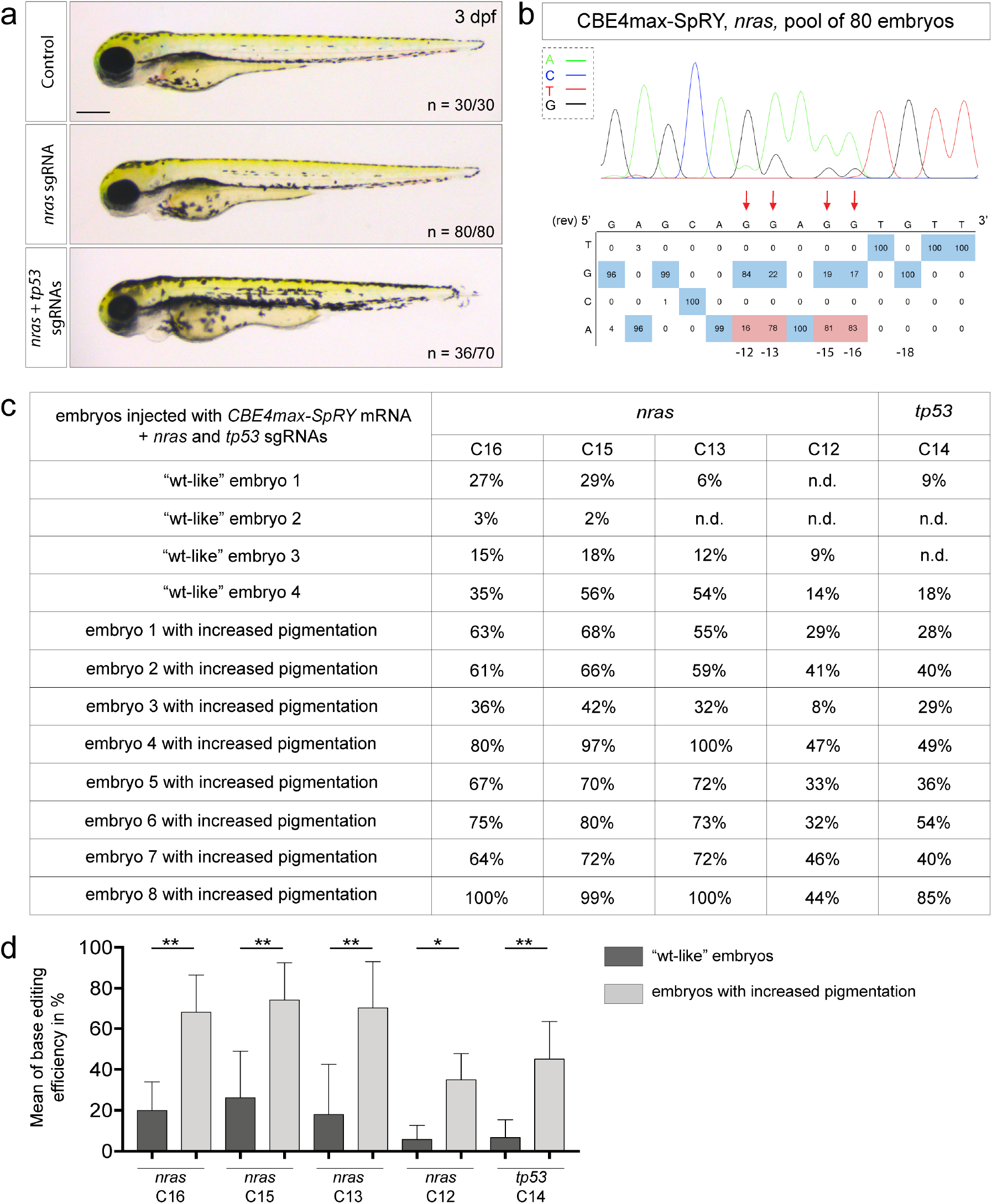
Endogenous activation of *nras* oncogene and knock-out of *tp53* tumor suppressor gene led to an increase of melanocyte numbers. **a**. Lateral view of 3 dpf embryos. The injected embryos edited only for *nras* do not present any defects whereas 50% of the injected embryos with *nras NA* and *tp53 Q170\** sgRNAs show an increase of pigmentation. Scale bar = 100 µm. **b**. DNA sequencing chromatogram of the targeted *nras* gene from a pool of 80 injected embryos with the *CBE4max-SpRY* mRNA and the *nras NA* sgRNA. C-to-T conversion shows 83% of efficiency for the C16, 81% for the C15, 78% for the C13 and 16% for the C12. Numbers in the boxes represent the percentage of each base at that sequence position. In red are highlighted the base substitutions introduced by base editing while the original bases are in blue. The color code of the chromatogram is indicated in the upper left corner (Adenine green, Cytosine blue, Thymine red, Guanine black). The distance from the PAM sequence of the targeted C base is indicated below the chromatogram^31^. **c**. C-to-T conversion efficiency for each targeted Cs in *nras* and *tp53* genes in single embryos injected with *CBE4max-SpRY* mRNA and *nras NA* and *tp53 Q170\** sgRNAs. 4 single embryos which did not show an increase of pigmentation named as “wt-like” embryos and 8 single embryos showing an increase of pigmentation have been sequenced. **d**. Histogram showing the mean with standard deviation of the base editing efficiency for the 4 “wt-like” embryos and the 8 embryos with increased pigmentation for Fig. 5c. Mann–Whitney test.

## Discussion

While the Base Editor technology is emerging as a revolutionary method to introduce precise single mutation in the genome, the presence at the good localization of the NGG PAM sequence is necessary and often a constrain. This has restricted its potential applications as the absence of the PAM puts the CBE unusable for many targets of interest. Here, we addressed these limitations in zebrafish by testing several base editor variants recognizing other PAM sequences. We unfortunately did not obtain any C-to-T conversions in zebrafish embryos using the previously published xCas9-BE4^20^ and CBE4max-SpG^14^. Nevertheless we could introduce point mutations with a remarkable high efficiency rate using the CBE4max-SpRY variant, a recently described near PAM-less CBE variant engineered and validated in cultured cells but never reported working in an animal model so far^14^. We thus significantly expand the base conversion possibilities in zebrafish and open the possibility to convert non NGG-targetable C bases which could not be achieved so far. Through our results, we could demonstrate that the CBE4max-SpRY can be extremely efficient, reaching 100% of C-to-T conversion in at least 50% of the injected embryos (Fig. 2d, e, 3b and 4c). We screened only 4 F0 fish and the 4 transmitted the mutation to the offspring with a high germline transmission rate (Supplementary Fig. 1). These last results are particularly remarkable as this efficiency rate has never been reached previously in zebrafish, even with the use of the classical CBEs recognizing the NGG PAM^9, 10, 11, 12^. Moreover, for one of our targets the CBE4max-SpRY was even more efficient than the ancBE4max (Fig. 2b-e). This could be due also to the fact that the CBE4max-SpRY might have a slightly different editing window that the usual PAM [-19, -13 bp] window as we show base editing for the C12 and no conversion for the C18 in *nras* targeting (Fig. 4). These last results and the PAM flexibility of the CBE4max-SpRY allow now to test several sgRNAs for the mutation of interest and play with the C base localization within the editing window to increase the C-to-T conversion efficacy compared to the use of the classical ancBE4max. These properties allow also to exclude other Cs present in the editing window to avoid the generation of other unwanted mutations near the targeted C. We furthermore demonstrated that using NA and NG PAMs we were able to precisely and simultaneously perform the Tyr (W273*) and Rb1 (W63*) mutations with high efficiency rates and both mutations were transmitted to the germline (Fig. 3). In this line, we also performed simultaneously 3 different and precise mutagenesis using 3 different PAM sequences (Fig. 4). This ability now in zebrafish is extremely useful if several mutations need to be introduced in order to model a human genetic disease such as cancers, especially if some mutations are located on the same chromosome. Different CRISPR co-selection methods have been engineered in *Drosophila, C. elegans* and cultured cells in order to phenotypically detect and enrich the cells or animals in which more mutagenesis events are taking place, by adding an sgRNA conferring a phenotypical read-out if the mutagenesis occurred^24, 25, 26, 27, 28^. Base editing co-selection strategies were recently demonstrated in cultured cells^21^ but have not been reported in animals so far. Moreover, these time-saving strategies have never been developed in zebrafish. We here developed a co-selection method for base editing in zebrafish to prescreen phenotypically injected embryos based on the detection of pigmentation defects. The method is based on the addition of the *tyrW273* NA* sgRNA to the micro-injection mix. Importantly mutations in the *tyr* gene are viable and do not affect most developmental processes in zebrafish. We indeed have shown in this study that using this strategy we could select the albino embryos which were the most C-to-T converted for *nras* and *rb1* genes (Fig. 4b, c). We also developed a zebrafish model combining activation mutation for *nras* oncogene and knock-out of *tp53* tumor suppressor gene revealing an increase of melanocytes (Fig. 5), a clear melanoma predisposing phenotype. Indeed, fish over-expressing human mutated HRAS oncogene in melanocytes were hyperpigmented at 3dpf and developed melanoma at the adult stage^15^. The high efficiencies of CBE4max-SpRY obtained in this study and the possibility to precisely mutate simultaneously several genes using different PAMs pave the way for future applications in a tissue specific manner and for genetic disease modeling. For example, it could be implemented in the MAZERATI (Modeling Approach in Zebrafish for Rapid Tumor Initiation) system^29^ in order to rapidly model and study *in vivo* combinations of endogenous mutations occurring in complex multigenic disorders. Finally, the high flexibility and efficiency of our method to induce combination of specific mutations will allow to rapidly create zebrafish cancer models combining the precise set of mutations found in individual patients. In the long term, these models could be used for rapid and patient specific drug screening for advanced personalized medicine^17, 30^.

## Acknowledgements

We thank Panagiotis Antoniou and Annarita Miccio for sharing the *pCAG-CBE4max-SpG-P2A-EGFP* and *pCAG-CBE4max-SpRY-P2A-EGFP* plasmids^14^. M.R. was supported by the Fondation pour la Recherche Médicale (FRM grant number ECO20170637481) and la Ligue Nationale Contre le Cancer. Work in the Del Bene laboratory was supported by ANR-18-CE16 “iReelAx”, UNADEV in partnership with ITMO NNP/AVIESAN (national alliance for life sciences and health) in the framework of research on vision and IHU FOReSIGHT [ANR-18-IAHU-0001] supported by French state funds managed by the Agence Nationale de la Recherche within the Investissements d’Avenir program.

## Author contributions

M.R. and M.S. did the experimental works and analyzed the results. M.C.M. helped the design of targets, reviewed and edited the manuscript. M.R., J.P.C., F.D.B. conceived the project and wrote the article. J.P.C., F.D.B. co supervised the study.

## Subjects

Prime editing, base editors, zebrafish, genome modifications

## Methods

### Fish lines and husbandry

Zebrafish (*Danio rerio*) were maintained at 28 °C on a 14 h light/10 h dark cycle. Fish were housed in the animal facility of our laboratory which was built according to the respective local animal welfare standards. All animal procedures were performed in accordance with French and European Union animal welfare guidelines. Animal handling and experimental procedures were approved by the Committee on ethics of animal experimentation.

### Molecular cloning

To generate the *pCS2+_CBE4max-SpG* and the *pCS2+_CBE4max-SpRY* plasmids, each *CBE4max-SpG* and *CBE4max-SpRY* sequence has been inserted into *pCS2+ plasmid* linearized with EcoRI using the Gibson Assembly Cloning Kit (New England Biolabs). The fragment has been amplified using the primers F-5’-TGCAGGATCCCATCGATTCGGCCACCATGAAACGGACAG -3’ and R-5’-TAGAGGCTCGAGAGGCCTTGTCAGACTTTCCTCTTCTTCTTGG -3’) from the *pCAG-CBE4max-SpG-P2A-EGFP* plasmid (Addgene plasmid #139998)^14^ and from the *pCAG-CBE4max-SpRY-P2A-EGFP* plasmid (Addgene plasmid #139999)^14^.

### mRNAs synthesis

*pCS2+_CBE4max-SpG* plasmid has been used to generate *CBE4max-SpG* mRNA *in vitro. pCS2+_CBE4max-SpRY* plasmid has been used to generate *CBE4max-SpRY* mRNA *in vitro*. Each plasmid was linearized with NotI restriction enzyme and mRNAs were synthesized by *in vitro* transcription with 1 µL of GTP from the kit added to the mix and lithium chloride precipitation (using the mMESSAGE mMACHINE sp6 Ultra kit #AM1340, Ambion).

### sgRNA design

The sequenceParser.py python script^12^ was used to design *tyr* sgRNAs. All the synthetic sgRNAs were synthesized by IDT as Alt-R® CRISPR-Cas9 crRNA. Prior injections, 2 µL of the crRNA (100 pmol/µL) and 2 µl of Alt-R® CRISPR-Cas9 tracrRNA (100 pmol/µL) from IDT were incubated at 95°C for 5 min, cooled down at room temperature and then kept in ice.

### Micro-injection

To make the mutagenesis with base editing, a mix of 1 nL of *CBE* mRNA and synthetic sgRNAs was injected into the cell at one-cell stage zebrafish embryos. For the single mutagenesis, the final concentration was 600 ng/μL for *CBE* mRNA and 43 pmol/μL for sgRNA. For the double *rb1* and *tyr* mutations, the final concentration was 600 ng/μL for *CBE* mRNA and 21 pmol/μL for each sgRNAs. For the double *rb1* and *nras* mutations, the final concentration was 600 ng/μL for *CBE* mRNA and 8,6 pmol/μL for *nras* sgRNA and 34,4 pmol/μL for *rb1* sgRNA. For the double *p53* and *nras* mutations, the final concentration was 600 ng/μL for *CBE* mRNA and 8,6 pmol/μL for *nras* sgRNA and 34,4 pmol/μL for t*p53* sgRNA. For the *tyr, nras* and *rb1* mutations, the final concentration was 600 ng/μL for *CBE* mRNA, 8,6 pmol/μL for *nras* sgRNA and 8,6 pmol/μL for *tyr* sgRNA and 25,8 pmol/μL for *rb1* sgRNA.

### Whole-embryos DNA sequencing

For genomic DNA extraction, embryos were digested for 1 h at 55°C in 0.5 mL lysis buffer (10 mM Tris, pH 8.0, 10 mM NaCl, 10 mM EDTA, and 2% SDS) with proteinase K (0.17 mg/mL, Roche Diagnostics) and inactivated 10 min at 95°C. To sequence and check for frequency of mutations, each target genomic locus was PCR-amplified using Phusion High-Fidelity DNA polymerase (Thermo Scientific). For the *tyr* locus, a PCR was performed with primers Fwd-5’-ATCGGGTGTATCTGCTGTTTTGG-3’ and Rev-5’-CCATACCGCCCCTAGAACTAACATT-3’. For the *rb1* locus, the primers used were Fwd-5’-TCTGTCAACTGTTGTTTTTCCAGAC-5’ and Rev-5’-CAATAAAAAGACAAGCTCCCCACTG-5’. For the *nras* locus, the primers used were Fwd-5’-CCTTTTCTCTCTTTTTGTCTGGGTG-5’ and Rev-5’-CGCAATCTCACGTTAATTGTAGTGT-5’. For the *tp53* locus, the primers used were Fwd-5’-ATATCTTGTCTGTTTTCTCCCTGCT-5’ and Rev-5’-GTCCTACAAAAAGGCTGTGACATAC-5’. The DNAs have been extracted on an agarose gel and purified (using the PCR clean-up gel extraction kit #740609.50, Macherey-Nagel) and the sanger sequencings have been performed by Eurofins. The sequences were analyzed using ApE software and quantifications of the mutation rates done using editR online software^31^.

### Imaging

Embryos were oriented in egg solution with an anesthetic (Tricaine 0,013%). Leica MZ10F was used to image them. Adult fish were imaged using a net and an Iphone xs.

### Statistics

A non-parametric t-test with the Mann–Whitney correction was applied to determine significance in base editing. The software used was Prism 7 (GraphPad).

## Supplementary information

### Disease modeling by efficient genome editing using a near PAM-less base editor *in vivo*

**Supplementary Figure 1.**
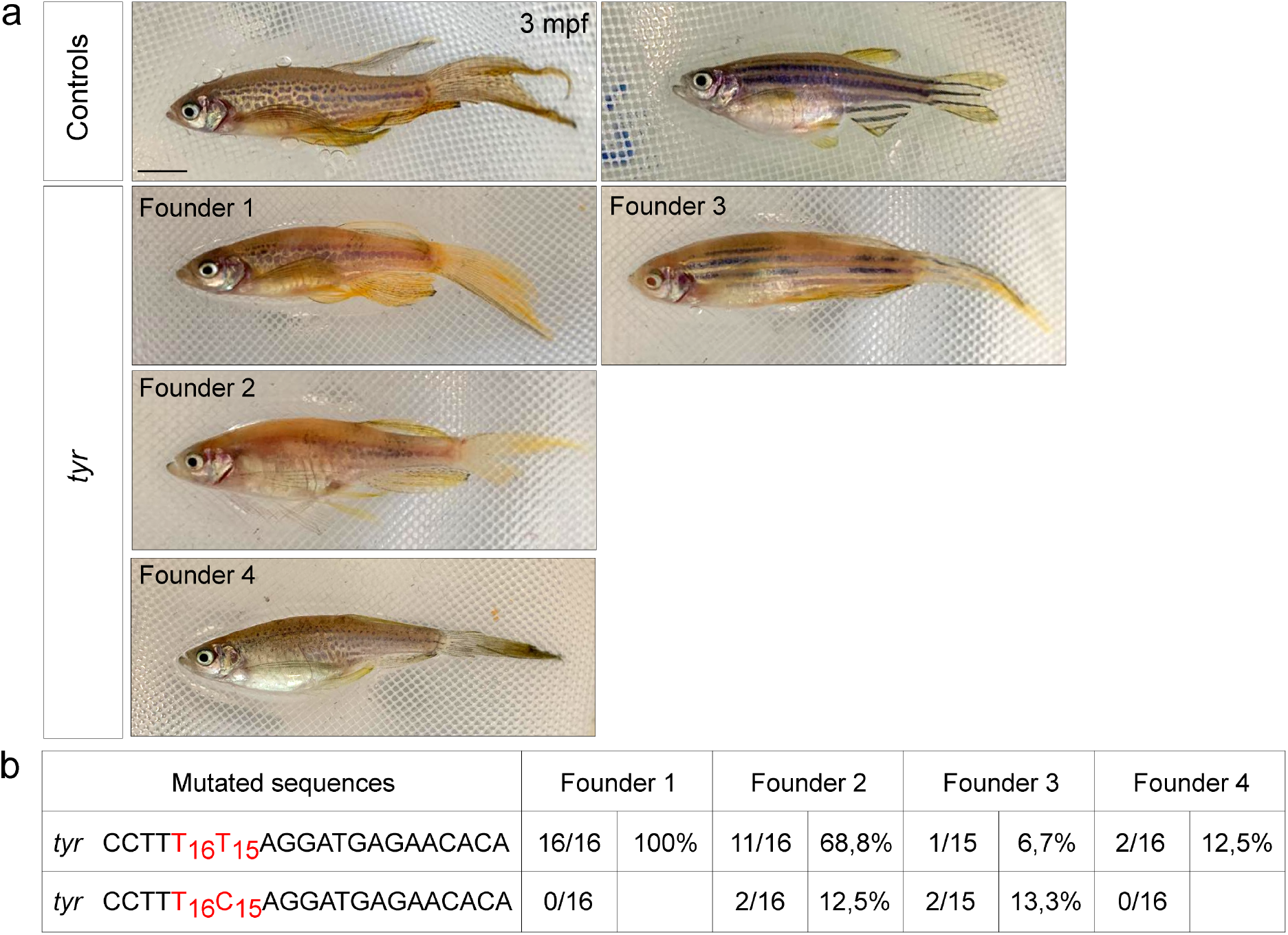
High germline transmission rate. **a**. Lateral view of a 3 months F0 fish injected at one-cell stage with the *CBE4max-SpRY* mRNA and the *tyrW273*NA* sgRNA and showing pigmentation lack defect. Scale bar = 5 mm. **b**. Sequenced *tyrosinase* locus of F1 single embryos randomly chosen from each founder. 16 embryos out of 16 were edited for *tyrosinase* for the Founder 1, 13 embryos out of 16 for the Founder 2, 3 embryos out of 15 for the Founder 3 and 2 embryos out of 16 for the Founder 4.

